# RNAhub2: prefiltering and compression enable scalable structure-based RNA discovery

**DOI:** 10.64898/2026.11.24.768864

**Authors:** Agata M. Kilar, William Gao, Nivedita Dutta, Marcin Magnus, Quentin Vicens, Elena Rivas

## Abstract

RNAhub is an established web server for structure-agnostic RNA homology search that builds multiple sequence alignments with nhmmer and performs downstream structural covariation analysis with R-scape and CaCoFold. RNAhub is efficiently applied to smaller genomes, including viruses, bacteria, and fungi. However, searches against large eukaryotic genomes remain computationally expensive, and genome storage requirements currently limit its taxonomic coverage, thereby reducing the opportunities to identify novel structural RNAs. Here, we present RNAhub2 which introduces three key advances to address those limitations: (i) query-adapted pre-filtering that creates a reduced query-specific search database prior to iterative alignment construction; (ii) parallelized genome-level searches with job resumption; (iii) Nucleotide Archival Format (NAF) compression of genomes, reducing storage by approximately 4.2-fold and enabling the addition of nearly 5,000 species across major eukaryotic kingdoms within the original storage quota without increasing computation time. These combined updates yield significantly improved runtimes and comparable or stronger covariation signal. RNAhub2 is now scalable for eukaryotic comparative RNA searches without sacrificing sensitivity. The web server is freely available at https://rnahub.org.

## INTRODUCTION

Many functional RNAs exert their biological roles through conserved RNA structures. An important signature of structural conservation is the presence of correlated substitutions (covariation) at paired positions in RNA sequence alignments, which preserve RNA base pairing (Westhof, 2026). Multiple sequence alignments provide a powerful approach for identifying novel conserved structural RNAs and for characterizing their secondary structures. Critically, the alignments used to infer putative structures must be constructed independently of any predicted structure to avoid circular reasoning (Gao et al., 2023).

RNAhub automates exploratory RNA analysis from a query sequence, by building structure-agnostic alignments which are evaluated by R-scape/CaCoFold (Rivas et al., 2017, 2020; Rivas, 2020) in order to identify evidence, or lack thereof, for the presence of a conserved RNA structure (Magnus et al., 2025).

The assessment of RNA structure by covariation analysis relies on the evolutionary information contained in the alignment. An informative alignment contains a balance between informative substitutions across related sequences, with sufficient diversity to detect covariation combined with enough conservation to preserve homology. An alignment with insufficient sequence variation may lack detectable covariation even when the RNA structure is conserved, whereas the absence of covariation in an alignment with adequate covariation-detection power provides evidence against a conserved structure (Rivas et al., 2020). The quality of this structural assessment strongly depends on the genome databases available to search. Access to more diverse genomes can improve RNAhub alignments by adding homologs with informative substitutions, increasing the power to detect covarying base pairs, and possibly enhancing structure prediction for RNA families. However, this is not automatic; more genomes may add sequences without variation that would increase redundancy and produce alignments that lack power (Rivas et al., 2020). Therefore, adding new genomes with taxonomic diversity enables broader taxonomic sampling, which would allow R-scape/CaCoFold to exploit a broader range of evolutionary variation. However, storing large genomes as uncompressed FASTA files takes substantial disc space, and the original RNAhub workflow was limited by storage capabilities, especially for plant and animal genomes.

The original RNAhub workflow was also limited by runtime. Genome-scale homology search is costly because it involves iterative searches with the nucleotide homology search tool nhmmer (Wheeler and Eddy, 2013). Iterative nhmmer searches across large eukaryotic genomes slow down web server interactions, especially in major eukaryotic kingdoms such as plants and animals. As public genome resources grow, these constraints hinder RNAhub’s ability to leverage evolutionary diversity.

Here we introduce RNAhub2, a redesigned workflow that overcomes both runtime and storage limitations while maintaining or improving the quality of the results. RNAhub2 is now available at https://rnahub.org/.

## RESULTS

### Accelerated homology searches by query-adapted database construction

The original RNAhub workflow builds alignments by iteratively searching the query sequence against a selected whole-genome database (Magnus et al., 2025). Although this strategy is effective, it requires iterative nhmmer (Wheeler and Eddy, 2013) searches over whole-genome collections, which makes searches against eukaryotic databases especially computationally expensive. To accelerate this step, RNAhub2 redesigns the workflow by introducing a query-specific prefiltering step before the original standard iterative RNAhub search (Fig. 1). In the RNAhub2 workflow, the query is first compared against the selected genome database using a lenient nhmmer E-value threshold (default E-value=10). Genomic regions detected in this broad search are all collected into a query-adapted database. The RNAhub2 iterative alignment-building procedure is then carried out against this reduced, query-dependent database rather than against the full genome collection used by RNAhub.

**Figure 1.**
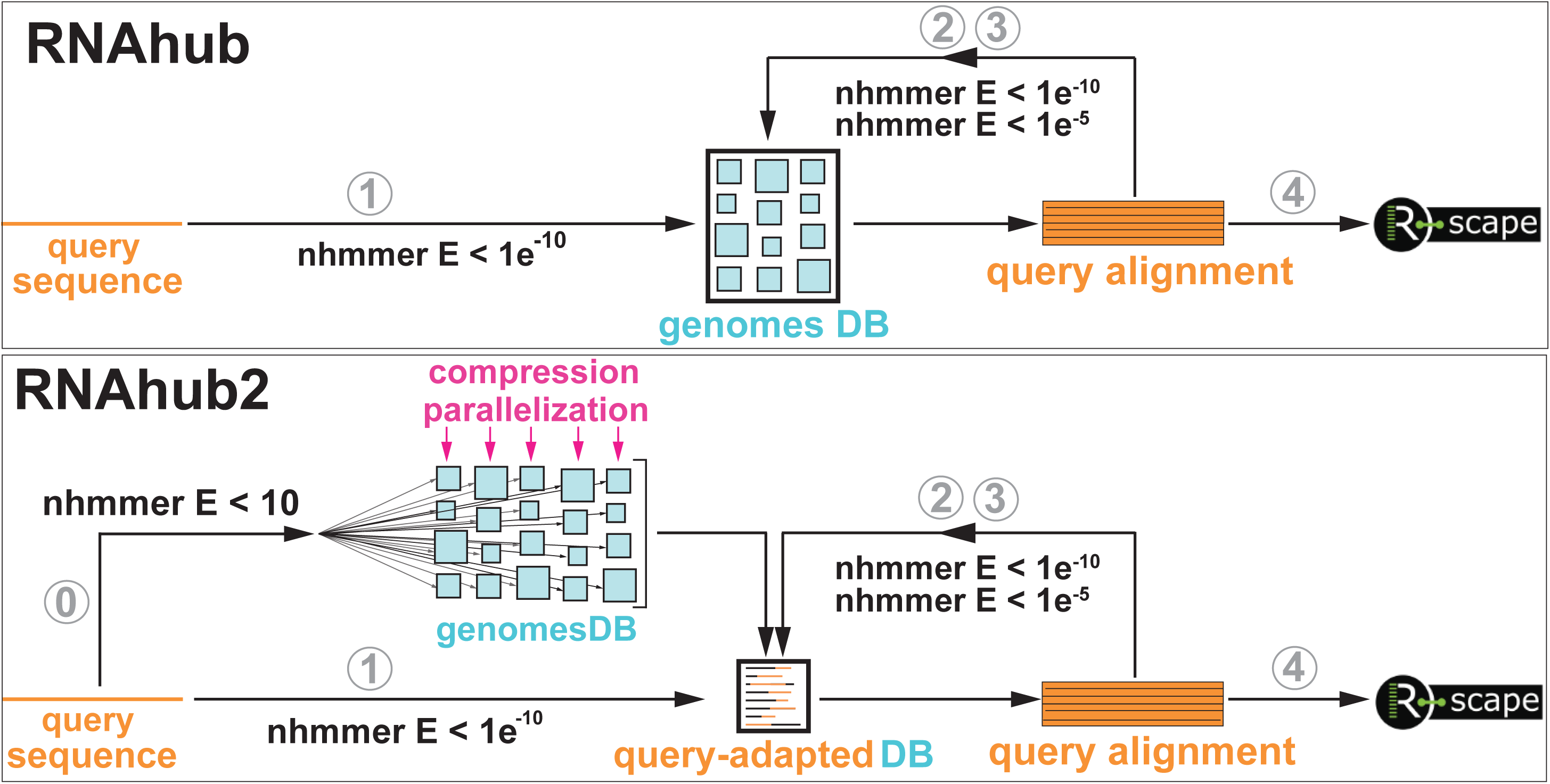
Comparison of the RNAhub and RNAhub2 workflows. Upper panel: In the RNAhub pipeline, the input query is compared against the full-genome database using nhmmer with custom E-value thresholds (1). The resulting hits are used to build a structure-agnostic multiple sequence alignment (2,3), which is then used for iterative model and alignment refinement and final structural evaluation with R-scape (4). Lower panel: In RNAhub2, an initial broad and lenient nhmmer search is performed in parallel across the database of individually compressed genomes to identify candidate regions and construct a query-adapted database (0). The original query is then searched against this reduced database (1). The resulting hits are used to build a structure-agnostic multiple sequence alignment, followed by iterative model and alignment refinement against the query-adapted database (2, 3). Each step uses the E-value thresholds of the corresponding step in RNAhub. The final alignment is passed to R-scape for downstream structural analysis (4).

This shift from the full genome database to a smaller, query-dependent database of possible homologs substantially reduces the search space used during iterative alignment construction. Instead of repeatedly searching gigabyte-scale genomic datasets, the RNAhub2 iterations are run on a much smaller database containing only candidate genomic regions. In this way, subsequent RNAhub2 alignment-building searches operate on a reduced dataset, decreasing runtime while preserving the overall RNAhub analysis strategy (Table 2).

To further improve performance, the RNAhub2 prefiltering step is parallelized by searching individual genomes separately rather than searching a single concatenated genome database. This also improves workflow robustness – if a search is interrupted, completed genome-level searches are retained and only unfinished searches need to be rerun, rather than restarting from the entire prefiltering step.

### NAF compression reduces genome storage with no increase in homology search time

The RNAhub webserver uses genome databases for homology searches and for building alignments of sequences homologous to a query. The rapidly growing number of genome assemblies, especially large eukaryotic genomes, creates an opportunity to expand the taxonomic diversity of the RNAhub database and to obtain more evolutionarily diverse alignments. However, adding more genomes greatly increases storage requirements, motivating the need for methods that reduce the size of individual genome files.

In order to address the genome storage limitation, we have tested different compression algorithms such as, `gzip`, `lz4`, `xz`, `zstd` and `NAF`. The goal is to maximally increase compression without sacrificing computational time necessary for the homology searches in the alignment building step. In our benchmark, `NAF` is highly efficient in both compression ratio and decompression time, for single genomes as well as for collections of imported databases (Kryukov et al., 2019). `NAF` achieves a storage compression factor of approximately 4.2 (Table 1, Supplemental materials, Fig. S1). Moreover, the nhmmer run time on `NAF`-compressed genomes is comparable to that on uncompressed genomes.

**Table 1.**
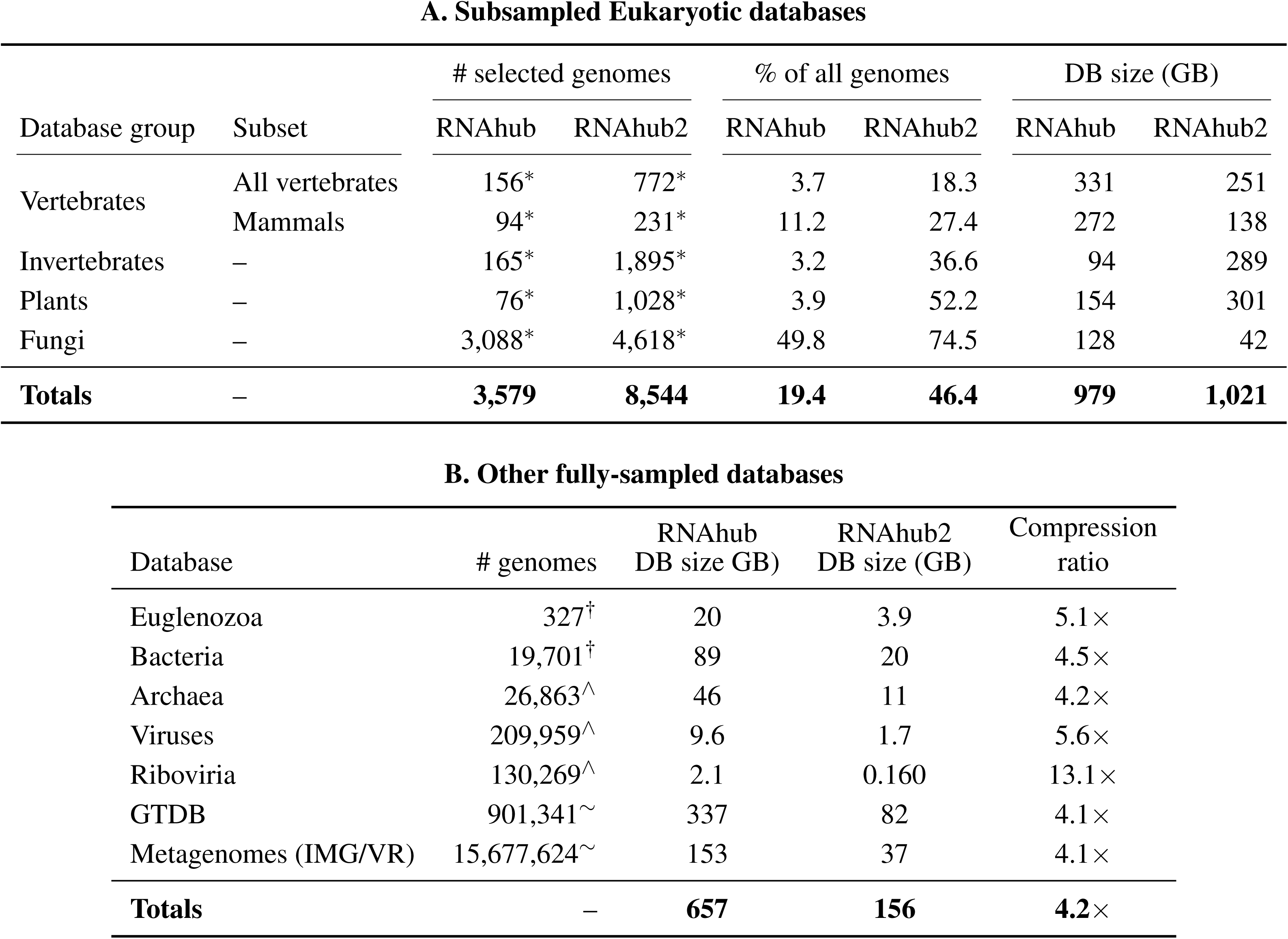
Comparison of the RNAhub and RNAhub2 genome databases. (A) Eukaryotic databases from which only a subset of the available genomes is included. (B) Effect of compression on other databases incorporated in full. The compression ratio is calculated as the RNAhub database size divided by the RNAhub2 database size. *^∗^* Database subsample; ^†^ All NCBI reference and representative genomes *^∧^* All NCBI genomes; *^∼^* All available databases.

| **A. Subsampled Eukaryotic databases** |  |  |  |  |  |  |  |
|---|---|---|---|---|---|---|---|
|  |  | # selected genomes |  | % of all genomes |  | DB size (GB) |  |
| Database group | Subset | RNAhub | RNAhub2 | RNAhub | RNAhub2 | RNAhub | RNAhub2 |
| Vertebrates | All vertebrates | 156<sup>*</sup> | 772<sup>*</sup> | 3.7 | 18.3 | 331 | 251 |
|  | Mammals | 94<sup>*</sup> | 231<sup>*</sup> | 11.2 | 27.4 | 272 | 138 |
| Invertebrates | – | 165<sup>*</sup> | 1,895<sup>*</sup> | 3.2 | 36.6 | 94 | 289 |
| Plants | – | 76<sup>*</sup> | 1,028<sup>*</sup> | 3.9 | 52.2 | 154 | 301 |
| Fungi | – | 3,088<sup>*</sup> | 4,618<sup>*</sup> | 49.8 | 74.5 | 128 | 42 |
| **Totals** | – | **3,579** | **8,544** | **19.4** | **46.4** | **979** | **1,021** |
| **B. Other fully-sampled databases** |  |  |  |  |  |  |  |
| Database | # genomes | RNAhub<br>DB size GB) | RNAhub2<br>DB size (GB) | Compression<br>ratio |  |  |  |
| Euglenozoa | 327<sup>†</sup> | 20 | 3.9 | 5.1× |  |  |  |
| Bacteria | 19,701<sup>†</sup> | 89 | 20 | 4.5× |  |  |  |
| Archaea | 26,863<sup>∧</sup> | 46 | 11 | 4.2× |  |  |  |
| Viruses | 209,959<sup>∧</sup> | 9.6 | 1.7 | 5.6× |  |  |  |
| Riboviria | 130,269<sup>∧</sup> | 2.1 | 0.160 | 13.1× |  |  |  |
| GTDB | 901,341<sup>∼</sup> | 337 | 82 | 4.1× |  |  |  |
| Metagenomes (IMG/VR) | 15,677,624<sup>∼</sup> | 153 | 37 | 4.1× |  |  |  |
| **Totals** | – | **657** | **156** | **4.2**× |  |  |  |

### RNAhub2 introduces a taxonomy-informed expansion of eukaryotic genomes

The original RNAhub genome-selection strategy did not explicitly account for taxonomic relationships. Genomes within a given structural cluster were treated as equivalent and selected at random, resulting in the preferential selection of genomes from taxonomic groups that were more abundant within that cluster (Magnus et al., 2025). For example, the RNAhub plant database contained only 4% of the available plant species, with 68 of the 76 selected genomes belonging to a single taxonomic class, Magnoliopsida (Fig 2 E).

**Figure 2.**
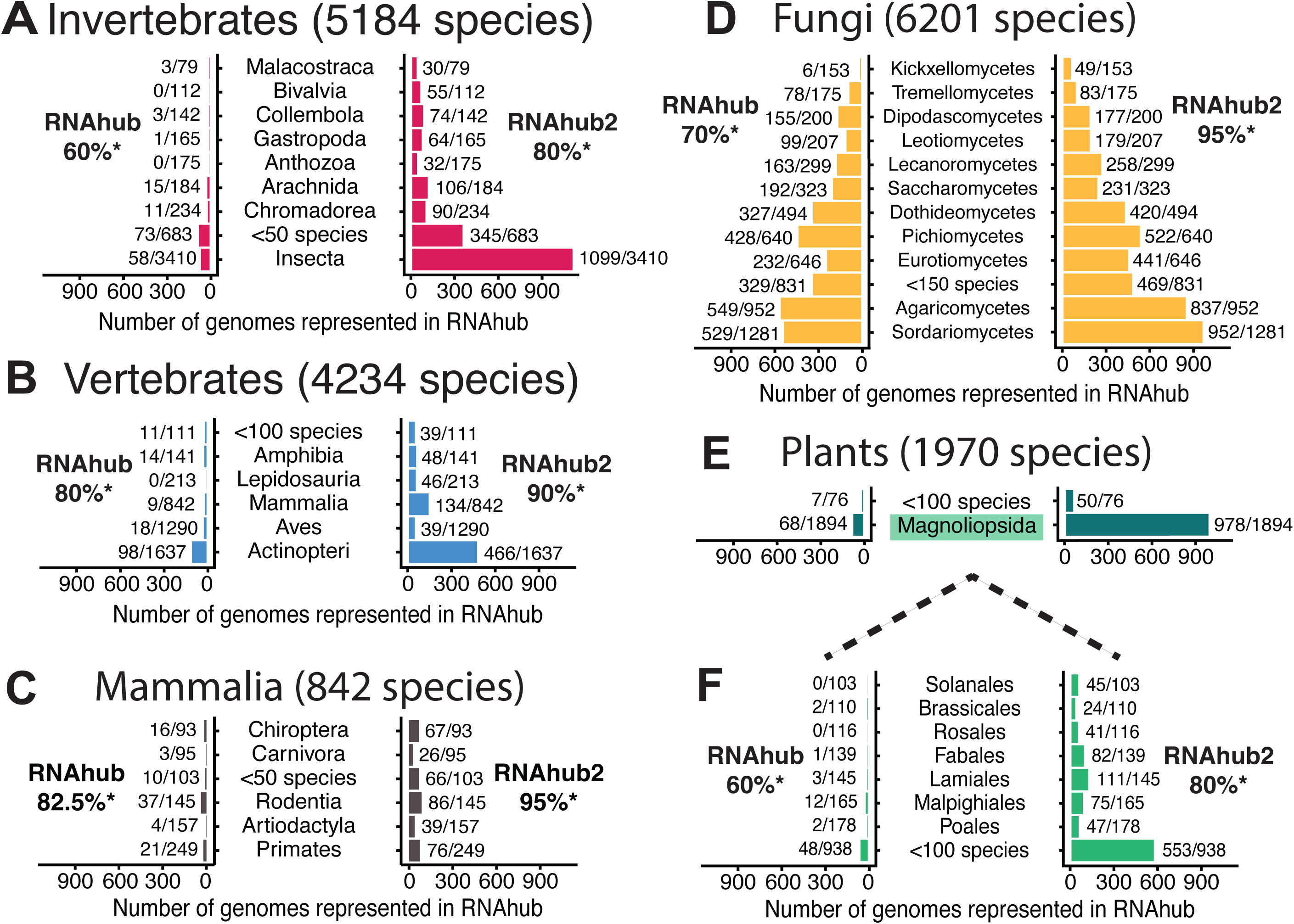
Eukaryotic taxonomic expansion in RNAhub2. Eukaryotic databases are downsampled due to storage constraints. Genomes were clustered by RNase MRP sequence identity, and only one representative per cluster was retained. A higher % identity clustering threshold produces more clusters and allows greater taxonomic diversity among the selected genomes. Because of genome compression, RNAhub2 is able to use higher % identity clustering thresholds than the original RNAhub, thereby retaining more genomes in all major eukaryotic groups: from 60% to 80% for (A) invertebrates, from 80% to 90% for (B) vertebrates, from 82.5% to 95% for (C) mammalia, from 70% to 95% for (D) fungi, and from 60% to 80% for (E) plants. (F) Magnoliopsida, the most highly represented plant class, is expanded to the order level to illustrate how its internal taxonomic coverage increases in RNAhub2. Mirrored bar plots show the number of genomes represented in RNAhub (left) and RNAhub2 (right) for the major taxonomic classes. Numbers next to the bars indicate the number of selected genomes relative to the total number available in each group; less populated classes were combined into the indicated aggregated categories.

In RNAhub2, genome compression enabled the inclusion of nearly 5,000 additional species while maintaining a comparable overall database size (Table 1A). A predefined storage limit was introduced for each eukaryotic database, allowing the number of included genomes to be increased until this limit was reached.

This approach increased genome representation across all major eukaryotic databases and enabled the inclusion of groups that had previously lacked representation, including corals and sea anemones (Anthozoa) and lizards, snakes, and tuatara (Lepidosauria). The largest increases were observed in the most species-rich groups: Insecta increased from 58 to 1,099 represented genomes, Magnoliopsida from 68 to 978, Actinopteri from 98 to 466, and Sordariomycetes from 529 to 952 (Fig. 2).

### RNAhub2 reduces runtime while preserving or improving covariation signal

To separate the contribution of query-adapted prefiltering from that of the expanded invertebrate genome database, we first benchmarked the effect of prefiltering alone. By default, the RNAhub uses the invertebrate-60 database (165 genomes), whereas RNAhub2 uses the expanded, invertebrate-80 database (1,895 genomes). For an exact comparison, we therefore also ran RNAhub2 on invertebrate-60 and compared both servers on eight annotated *C. elegans* non-coding RNAs. RNAhub searches uncompressed genomes, while RNAhub2 operates only on compressed genomes. However, the search tool used in the prefiltering step has a similar runtime on compressed and uncompressed genomes (Supp. Fig. S1), so compression does not account for the observed differences.

As described in Table 2, on the invertebrate-60 database, RNAhub2 reduced runtime substantially for every query, in several cases by more than 10-fold (e.g. RNase P: from 94 to 8 min; U1: from 82 to 7 min), with a smaller speed up (3-fold time decrease) for SSU (from 267 down to 88 min). Sequence counts, average pairwise identity, and observed covarying base pairs were similar between RNAhub and RNAhub2 at those matched database. The exceptions are RNase P and C/D-box snoRNA, where RNAhub2 recovered more homologs (RNase P: 21 vs 4; C/D-box: 59 vs 3), with lower pairwise identity in both cases.

**Table 2.**
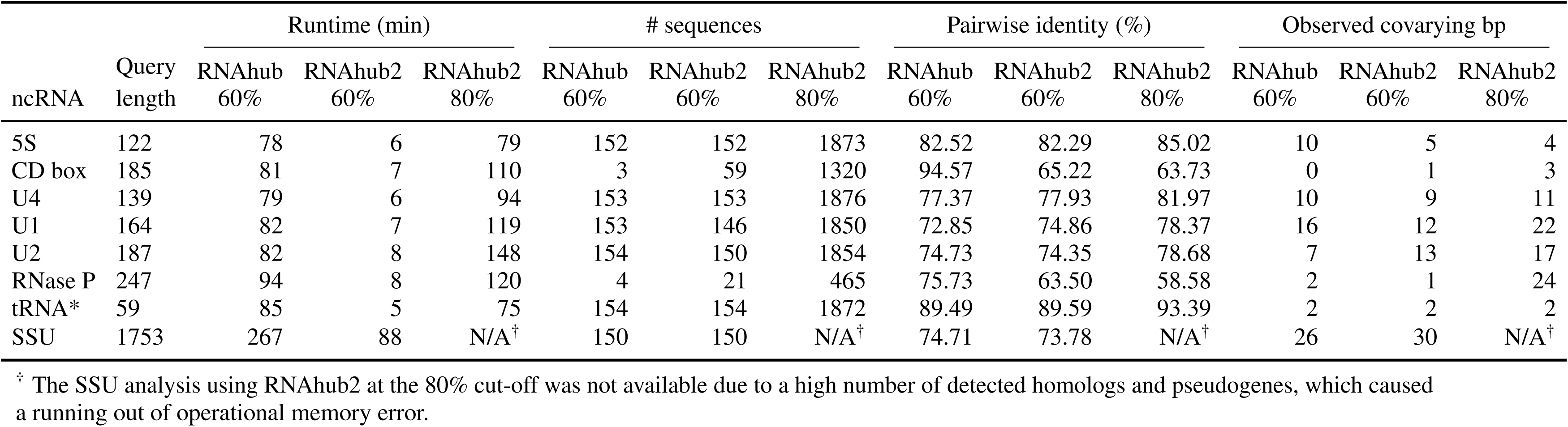
Runtime and alignment statistics for RNAhub and RNAhub2. Column labels give the tool and the clustering threshold used to build database, with 60% and 80% denoting invertebrate-60 database (165 genomes) and invertebrate-80 (1,895 genomes)

|  |  | Runtime (min) |  |  | # sequences |  |  | Pairwise identity (%) |  |  | Observed covarying bp |  |  |
|---|---|---|---|---|---|---|---|---|---|---|---|---|---|
| ncRNA | Query<br>length | RNAhub<br>60% | RNAhub2<br>60% | RNAhub2<br>80% | RNAhub<br>60% | RNAhub2<br>60% | RNAhub2<br>80% | RNAhub<br>60% | RNAhub2<br>60% | RNAhub2<br>80% | RNAhub<br>60% | RNAhub2<br>60% | RNAhub2<br>80% |
| 5S | 122 | 78 | 6 | 79 | 152 | 152 | 1873 | 82.52 | 82.29 | 85.02 | 10 | 5 | 4 |
| CD box | 185 | 81 | 7 | 110 | 3 | 59 | 1320 | 94.57 | 65.22 | 63.73 | 0 | 1 | 3 |
| U4 | 139 | 79 | 6 | 94 | 153 | 153 | 1876 | 77.37 | 77.93 | 81.97 | 10 | 9 | 11 |
| U1 | 164 | 82 | 7 | 119 | 153 | 146 | 1850 | 72.85 | 74.86 | 78.37 | 16 | 12 | 22 |
| U2 | 187 | 82 | 8 | 148 | 154 | 150 | 1854 | 74.73 | 74.35 | 78.68 | 7 | 13 | 17 |
| RNase P | 247 | 94 | 8 | 120 | 4 | 21 | 465 | 75.73 | 63.50 | 58.58 | 2 | 1 | 24 |
| tRNA* | 59 | 85 | 5 | 75 | 154 | 154 | 1872 | 89.49 | 89.59 | 93.39 | 2 | 2 | 2 |
| SSU | 1753 | 267 | 88 | N/A<sup>†</sup> | 150 | 150 | N/A<sup>†</sup> | 74.71 | 73.78 | N/A<sup>†</sup> | 26 | 30 | N/A<sup>†</sup> |

<sup>†</sup> The SSU analysis using RNAhub2 at the 80% cut-off was not available due to a high number of detected homologs and pseudogenes, which caused a running out of operational memory error.

The computational gain from query-adapted prefiltering can also be viewed as enabling database expansion rather than simply shortening the time of individual searches. Although the invertebrate-80 database contains approximately 11.5-fold more genomes than invertebrate-60, RNAhub2 searches against the expanded database generally completed on a timescale comparable to RNAhub searches against the original database.

To asses the effect of extending genome representation, we compared RNAhub on the default invertebrate-60 database, and RNAhub2 on the expanded invertebrate-80 database (Table 2). Both databases were constructed using the RNase MRP-based clustering method described in (Magnus et al., 2025), retaining one representative genome per cluster at sequence identity thresholds of 60% and 80%, respectively. Higher thresholds produce more clusters, resulting in a broader genome representation in the invertebrate-80 database with respect to invertebrate-60.

Table 2 shows that searching against invertebrate-80 further increased alignment depth for every query, increased observed covarying base pairs for most queries, notably RNase P, U1, U2, and C/D box, with 5S as the exception. Average sequence identity of the alignment decreased further for RNase P and C/D-box, and increased slightly for the rest. SSU ran out of memory at invertebrate-80 due to a large number of gene copies and pseudogenes detected.

Figure 3 shows all these per-query trends for all eight ncRNA tests by plotting alignment power against observed significant covariation. With RNAhub (Fig. 3A left), most queries had low statistical power to detect covariation and even lower observed covariation, with RNase P and the C/D-box snoRNA falling close to the 0. On the other hand, with RNAhub2 (Fig. 3A right), the same queries shifted to higher power and higher observed covariation, with RNase P and U1 reaching the highest values in the panel. Each structural RNA remained below the diagonal, showing that additional power was not fully converted into significant covariation, but the gain in both power and observed covariation was consistent across nearly all queries.

**Figure 3.**
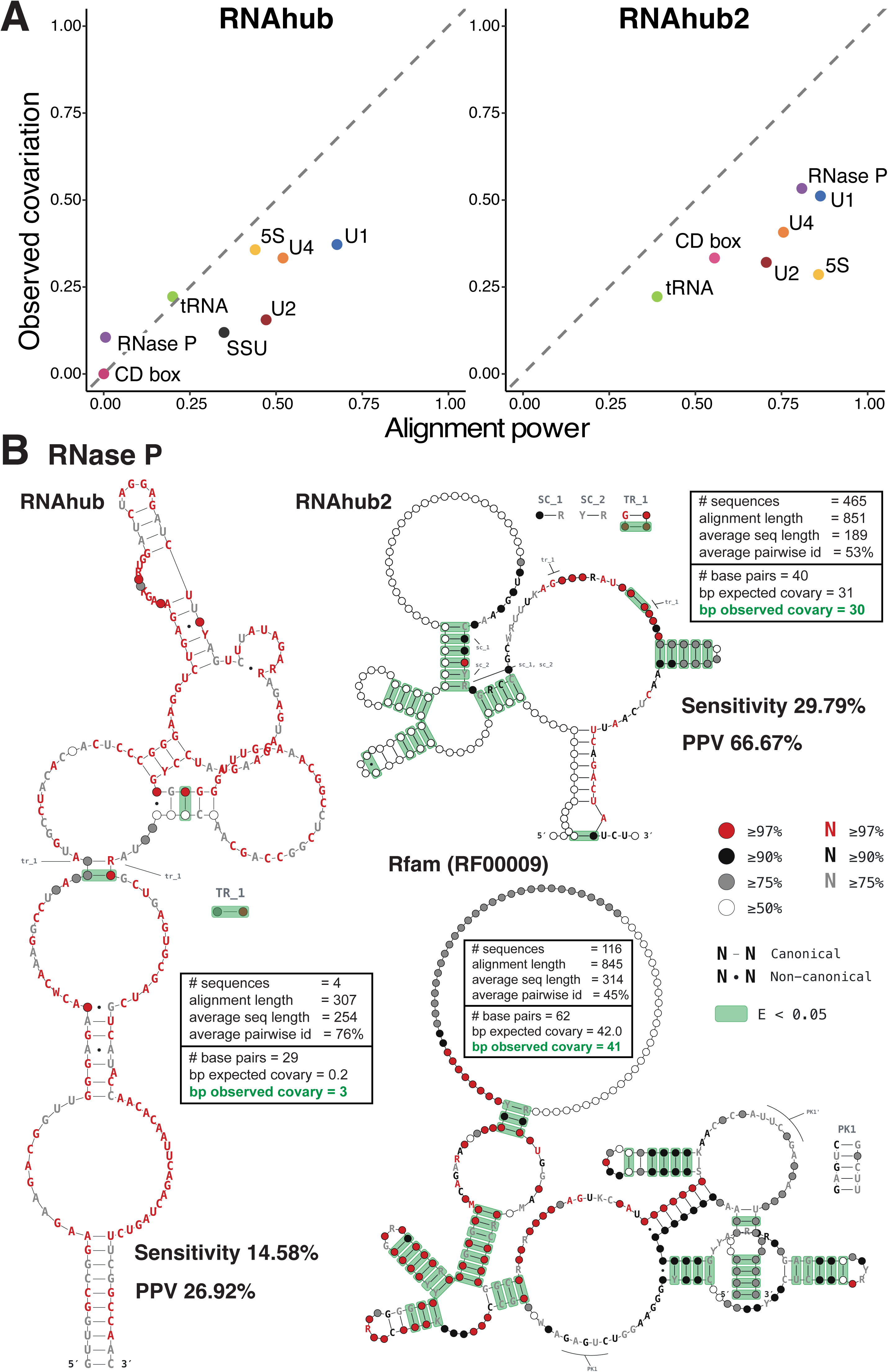
Benchmark of RNAhub and RNAhub2 implementations. A) Comparison of power and significant covariation in *C. elegans* ncRNA alignments obtained with RNAhub and RNAhub2, with Invertebrates databases, containing 165 and 1,895 genomes, respectively. Alignment power is shown as the number of expected covarying base pairs divided by the total number of base pairs in the consensus structure. Observed covariation is shown as the number of significantly covarying base pairs divided by the total number of base pairs. The dashed line indicates equal proportions of expected and observed covarying base pairs. Each point represents an ncRNA, with matching colors identifying the same ncRNA in the RNAhub and RNAhub2 plots. The complete set of tested sequences and alignment statistics is provided in Table 2 and described in Methods section. B) RNase P as an illustrative example of the benchmark outcome. Structures were plotted using RFview (Incarnato et al., 2018). Alignment and R-scape statistics are shown in the boxes. Sensitivity and positive predicted value (PPV) of all predicted base pairs were calculated relative to the Rfam family structure (RF00009) using `esl-compstruct` from HMMER suite.

RNase P RNA illustrates this improvement in detail (Fig. 3B). The original RNAhub alignment, built from only 4 sequences at 76% average pairwise identity, contained 29 predicted base pairs, of which 0.2 were expected to covary by chance. Nonetheless, three covarying base pairs reached significance. Comparing all 29 predicted base pairs of this alignment against the curated Rfam reference structure (RF00009) showed low agreement (sensitivity 14.58%, PPV 26.92%). The RNAhub2 alignment, built from 465 sequences at 53% average pairwise identity, contained 40 predicted base pairs with 31 expected to covary, of which 30 reached significance. Comparing all 40 predicted base pairs against the same reference structure (116 sequences, 45% average identity, 62 base pairs, 41 of which were found to covary significantly) showed markedly higher agreement, with sensitivity increasing from 14.58% to 29.79% and PPV from 26.92% to 66.67% relative to RNAhub.

### Expanded vertebrate genome database strengthens covariation support for telomerase RNA

Telomerase RNA (TR) exhibits remarkable evolutionary diversity in sequence, length, and biogenesis across eukaryotes (Fajkus et al., 2021, 2023). Within vertebrates, substantial primary-sequence divergence coexists with conserved structural domains required for telomerase function (Chen et al., 2000; Seemann et al., 2022). This combination makes TR a challenging example for automated homolog recruitment and alignment. We therefore compared RNAhub and RNAhub2 using human TR as the query to test whether expanded vertebrate genome sampling strengthens comparative structural evidence.

Using RNAhub, we obtained an alignment of 56 sequences with an average pairwise identity of 53%; R-scape detected one significant covarying base pair (Fig. 4). RNAhub2, using the expanded vertebrate database, yielded an alignment of 290 sequences with an average identity of 62% and detected five significant covarying pairs across four stems, while approximately halving the computational time.

**Figure 4.**
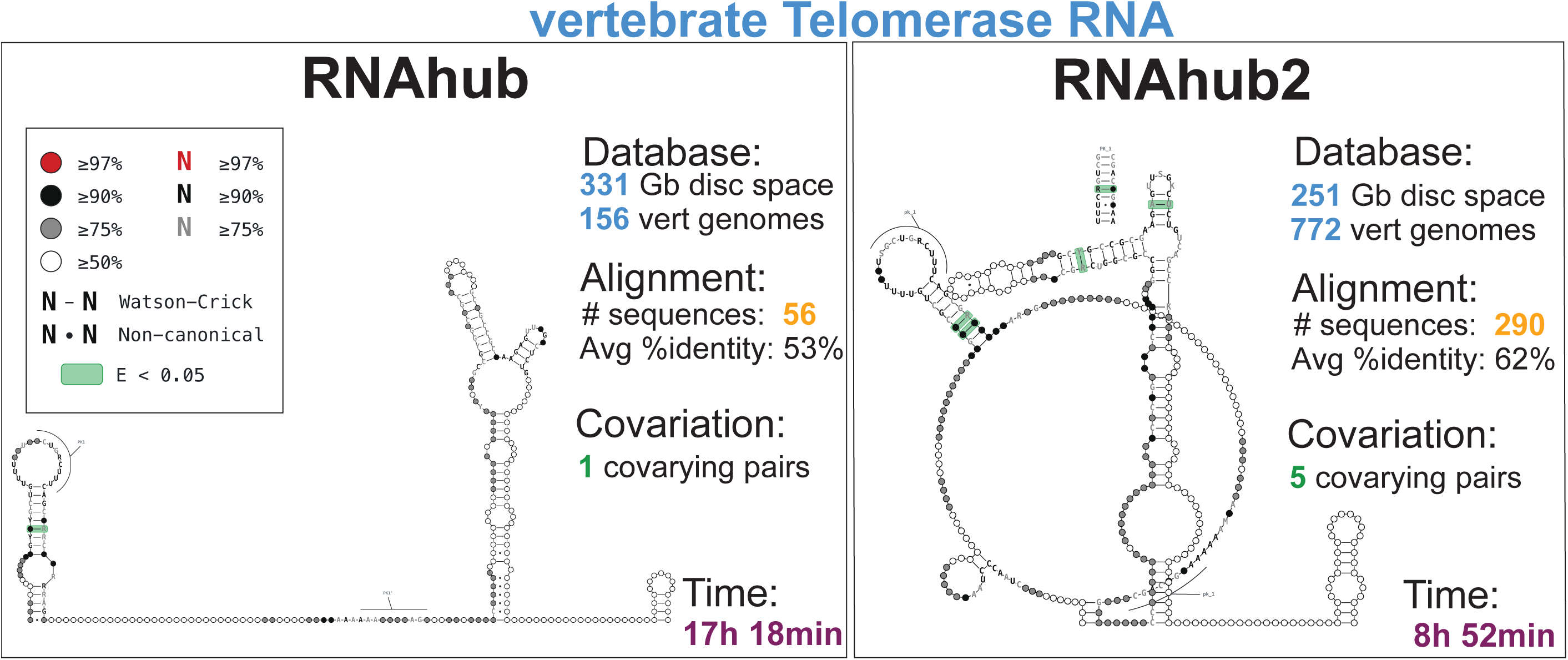
Comparison of RNAhub and RNAhub2 using human telomerase RNA. Human telomerase RNA (TR) was used as the query to search the original Vertebrates database with RNAhub (left) and the expanded Vertebrates database with RNAhub2 (right). Each panel reports the database storage requirement and number of genomes, the number of aligned sequences and mean pairwise identity, the number of significantly covarying base pairs, and the analysis runtime. Covariation detected by R-scape in the final sequence-based alignments is displayed on the telomerase secondary structure using RFview (Incarnato et al., 2018). Green shading marks significantly covarying pairs (*E <* 0.05). Nucleotide colors indicate sequence conservation, and lines and dots denote Watson–Crick and non-canonical pairs, respectively, as defined in the legend. Matching PK labels identify the paired regions forming pseudoknots.

The increased covariation signal despite higher average sequence identity illustrates that stronger structural evidence need not require a more divergent alignment on average. Additional homologs can contribute informative substitutions even when average identity increases. These results are consistent with a benefit from expanded vertebrate sampling and demonstrate improved recovery of covariation support for a sequence-divergent RNA.

### CASP benchmarking further highlights gains from expanded eukaryotic coverage and prefiltering

CASP provides experimentally characterized RNA targets collected for blind structure prediction, offering an external benchmark for comparing homolog recruitment and covariation support between RNAhub and RNAhub2. Among the CASP targets considered for this benchmark, three were of eukaryotic origin and suitable for RNAhub analysis: the CASP16 target R1260, a group I intron from *Tetrahymena*; the CASP15 target R1107, the human CPEB3 ribozyme; and the CASP17 target R2415s0, a U1 snRNA from an unidentified organism.

For CASP16 R1260, RNAhub detected no homologs and stopped after the first iteration because no sequence met the inclusion E-value threshold, leaving only the original query, while RNAhub2 recruited 20 sequences from the expanded Invertebrates database (Fig. 5A). Only one species represented in the final RNAhub2 alignment, *Geonemertes pelaensis* (`GCA 041107555.1`), was present in the original RNAhub database. The *G. pelaensis* candidate was not recruited during the first iteration but entered the alignment during the second iteration, after homologs from the expanded database had contributed to the search profile. The RNAhub2 third-round alignment contained 7 significantly covarying pairs and 25 after realignment based on structure with Infernal (Nawrocki and Eddy, 2013). The final CaCoFold model contained 74 base pairs, of which 18.1 *±* 3.3 were expected to covary. Comparison with the experimentally determined structure was restricted to target positions 69–309, corresponding to the region covered by the RNAhub2 model, and yielded a sensitivity of 58% and a positive predictive value of 76%. The model includes the P5b stem–loop, with four significantly covarying stem pairs and a conserved GAAA tetraloop. In the experimental *Tetrahymena* ribozyme structure, this tetraloop contacts a receptor in the P6 region (Su et al., 2021). The comparative analysis supports the local P5b stem–loop but does not test or establish this tertiary interaction.

**Figure 5.**
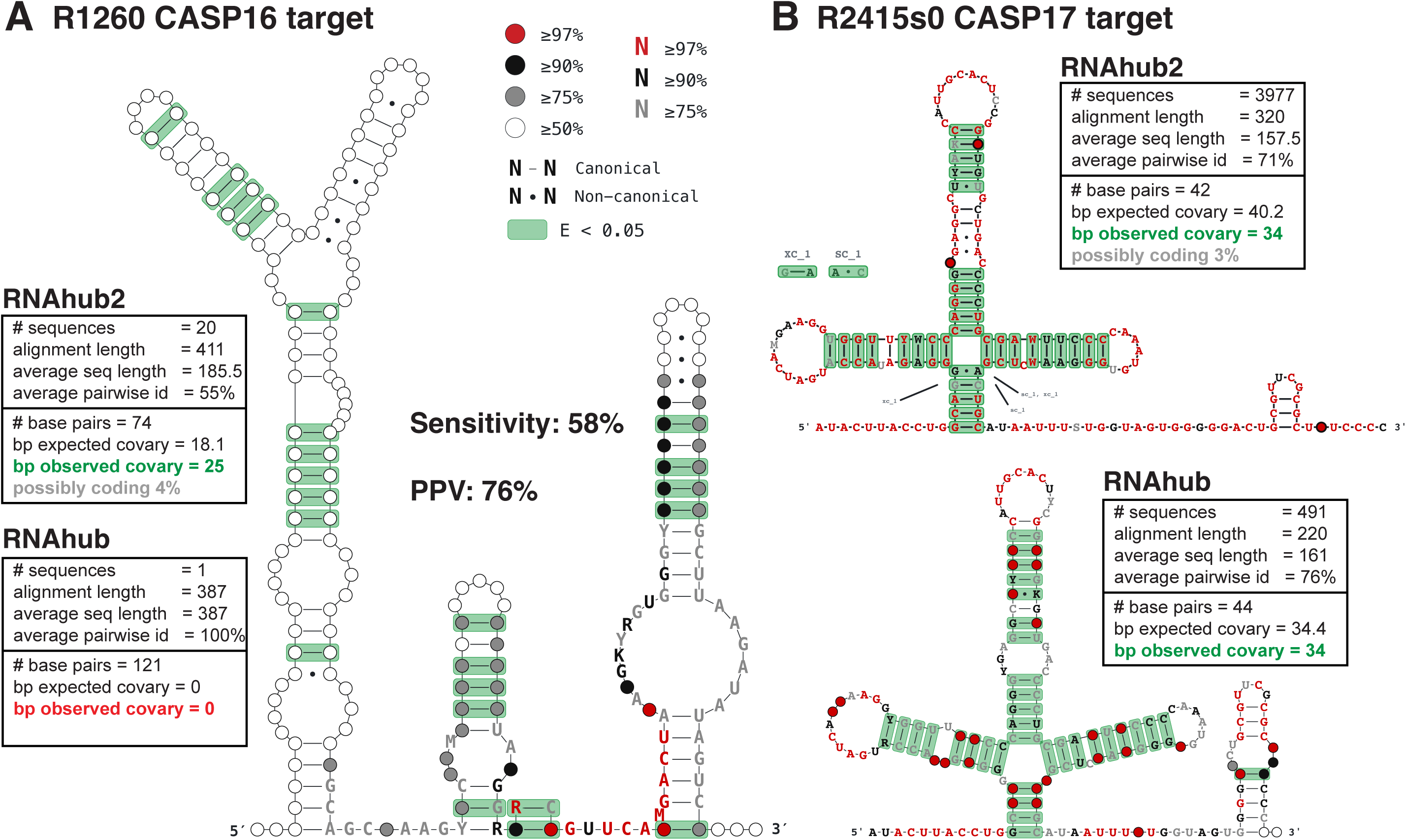
Comparison of RNAhub and RNAhub2 on selected eukaryotic CASP targets. RNAhub was searched against the original genome databases, and RNAhub2 against the corresponding expanded databases. Boxes report the number of sequences, alignment length, mean sequence length, mean pairwise identity, number of base pairs in the CaCoFold model, and expected and observed numbers of significantly covarying base pairs. Statistics are shown after Infernal realignment. A) P4-P6 region from CASP16 target R1260, a group I intron from *Tetrahymena*, was searched against the respective Invertebrates databases. The RNAhub2 structure was rendered with RFview (Incarnato et al., 2018). Green shading marks significantly covarying pairs (*E <* 0.05), and nucleotide colors indicate sequence conservation. Because the RNAhub2 model covers only positions 69–309 of the 387-nt target, sensitivity and positive predictive value (PPV) were calculated only for this fragment. Canonical base pairs were compared with the corresponding region of the experimental structure from PDB chain `9CBU N` using `esl-compstruct` from the HMMER suite. B) CASP17 target R2415s0, a U1 snRNA from an unidentified organism, was searched against the combined Mammals, Vertebrates, Invertebrates, Fungi, and Plants databases.

For CASP17 R2415s0, searches against the combined Mammals, Vertebrates, Invertebrates, Fungi, and Plants databases produced alignments of 491 sequences with RNAhub and 3,977 sequences with RNAhub2. After Infernal realignment, both analyses detected 34 significantly covarying pairs. The RNAhub model contained 44 proposed base pairs, with 34.4 expected to covary, whereas the RNAhub2 model contained 42 proposed base pairs, with 40.2 expected to covary.

For CASP15 R1107, searches against the combined Mammals and Vertebrates databases produced alignments of 127 sequences with RNAhub and 309 sequences with RNAhub2. The respective CaCoFold models contained 18 and 17 proposed base pairs, with 6.4 and 8.2 pairs expected to covary; neither analysis detected a significantly covarying pair, and Infernal realignment was therefore not performed.

Together, these examples show that RNAhub2 increased homolog recruitment across the three selected eukaryotic targets, but the effect on covariation support varied: significant covariation became detectable for R1260, remained undetected for R1107, and was recovered at the same level for R2415s0.

Comparative performance of RNAhub and RNAhub2 in a selected subset of other eukaryotic structural RNAs is provided in Supplemental Table S1.

### Analysis of CASP16 bacterial and viral targets

The bacterial and viral genome collections were unchanged between RNAhub and RNAhub2, allowing us to examine how prefiltering affects homolog recruitment and covariation support without the accompanying expansion of these databases. Table 3 compares the two workflows using a selected set of 11 CASP16 RNA targets representing several established structural RNA families, including catalytic introns, a riboswitch, and other structured non-coding RNAs. The range of alignment depths and sequence identities represented in this set provides different contexts in which to assess the retention of structural evidence.

**Table 3.** Alignment statistics for selected viral and bacterial CASP16 RNA targets analyzed in unflanked mode. Statistics for the final nhmmer alignment (v3 rm) are shown in black (top), and those for the Infernal realignment are shown in blue (bottom). Numbers in parentheses indicate significantly covarying pairs classified by R-scape as possibly arising from protein-coding constraints.

| CASP ID<br>Rfam ID | RNA class | Database | Pairwise ID |  | #sequences |  | #covarying pairs |  |
|---|---|---|---|---|---|---|---|---|
|  |  |  | RNAhub | RNAhub2 | RNAhub | RNAhub2 | RNAhub | RNAhub2 |
| R1211<br>RF00386 | CVB3 cloverleaf RNA<br>in complex with 3C protease | Viral | 82.7%<br>82.6% | 81.62%<br>81.79% | 48<br>48 | 48<br>48 | 1<br>8 | 2<br>9 |
| R1241<br>RF02001 | Group II intron | Bacteria | 52.13%<br>53.36% | 53.44%<br>55.48% | 3197<br>3197 | 2673<br>2673 | 124 (31c)<br>147 (37c) | 108 (21c)<br>128 (25c) |
| R1242<br>RF03072 | RaiA RNA motif | Bacteria+GTDB | 68.48%<br>68.3% | 69.55%<br>67.34% | 1034<br>1034 | 1160<br>1160 | 34 (4c)<br>47 (5c) | 22<br>26 |
| R1289<br>RF00028 | Precursor tRNA | Bacteria+GTDB | 58.71%<br>58.21% | 60.34%<br>59.65% | 1903<br>1903 | 2082<br>2082 | 139 (56c)<br>156 (75c) | 90 (19c)<br>109 (28c) |
| R1261<br>RF01750 | ZTP Riboswitch | Bacteria+GTDB | 83.42%<br>82.58% | 82.53%<br>82.42% | 161<br>161 | 167<br>167 | 1<br>2 | 1<br>2 |
| R0252<br>RF03087 | ROOL ncRNA<br>(ucc) | Bacteria+GTDB+IMGVR | 51.41%<br>51.17% | 62.83%<br>62.41% | 163<br>163 | 42<br>42 | 11 (1c)<br>36 (1c) | 1<br>20 |
| R0253<br>RF03087 | ROOL ncRNA<br>(env) | Bacteria+GTDB+IMGVR | 50.56%<br>49.65% | 53.12%<br>51.14% | 3352<br>3352 | 885<br>885 | 130 (17c)<br>183 (30c) | 73 (2c)<br>113 (14c) |
| R0254<br>RF02032 | GOLLD 3’ domain | Bacteria+GTDB+IMGVR | 59.13%<br>58.38% | 62.98%<br>62.68% | 11503<br>11503 | 9110<br>9110 | 122 (24c)<br>143 (37c) | 98 (18c)<br>119(17c) |
| R0283<br>RF03087 | Enterococcus ncRNA | Bacteria | 50.99%<br>50.14% | 53.35%<br>48.86% | 31<br>31 | 23<br>23 | 4<br>21 | 2<br>17 |
| R0285<br>RF01071 | OLE | Bacteria | 67.02%<br>66.77% | 66.96%<br>66.75% | 804<br>804 | 803<br>803 | 72<br>105 | 66<br>108 |
| R1290<br>RF02001 | Group II intron<br>HYER1 without TRS | Bacteria | 48.06%<br>50.8% | 46.4%<br>51.85% | 3239<br>3239 | 2792<br>2792 | 151 (40c)<br>273 (127c) | 117 (24c)<br>196 (62c) |

Both workflows recovered significant covariation for all tested targets, although RNAhub2 yielded fewer significant pairs in several cases. ROOL R0252 illustrates an interesting case: RNAhub2 recovered 42 sequences with 20 significant covarying pairs, compared with 163 sequences and 36 significant covarying pairs using RNAhub. The second ROOL target available in CASP16 benchmark, R0253, produced a substantially better outcome: RNAhub2 recovered 885 sequences and detected 113 significant covarying pairs, while RNAhub recovered 3352 sequences and detected 183 significant covarying pairs. The CASP16 organisers likewise reported a limited homologs recruitment for R0252 target, whose rMSA alignment was approximately 12-fold shallower than those of other targets (Kretsch et al., 2026). These observations suggest that limited homolog recruitment may constrain covariation detection for R0252.

The differences between workflows were target-dependent and were not explained by alignment depth alone. RNAhub2 recruited more sequences for RaiA and the precursor tRNA target, yet recovered fewer significantly covarying pairs after Infernal realignment. Conversely, for OLE, RNAhub2 recovered 108 significant pairs, while RNAhub recovered 105 significant pairs with almost identical alignment depths. Infernal realignment also substantially increased the detected covariation for several targets without changing the reported sequence counts, highlighting the contribution of the alignment stage to the observed results. These comparisons indicate that homolog recruitment and the resulting alignment both need to be considered when assessing the retention of comparative structural evidence.

Overall, RNAhub2 retained detectable covariation for all 11 selected CASP16 targets, although the extent of this support varied between workflows and was not explained by alignment depth alone.

### Analysis and visualization improvements

We added several features to RNAhub2 that make searches more flexible and results easier to interpret. All described below updates are also available for RNAhub.

Users can now start a search from an alignment instead of a single query sequence. RNAhub2 accepts unaligned FASTA and Stockholm files, and checks the upload before the job is created. The uploaded alignment is then used as the query input for the iterative search.

When a job is submitted, the query, or the first sequence of an uploaded alignment, is also screened with tRNAscan-SE 2.0 (Chan et al., 2021). Any tRNA genes it predicts are reported with their isotype, anticodon and cloverleaf secondary structure, so users can recognize tRNA or tRNA-derived queries before interpreting the homology search. Significant hits, marked “!” by Infernal, are shown in full, and hits below the inclusion threshold are shown faded.

RNAhub2 also checks whether an experimentally determined 3D structure is available for the query or its homologs. We built a nonredundant database of all nucleic-acid chains in the Protein Data Bank (Berman et al., 2000) from the wwPDB sequence file, collapsing identical sequences into a single entry. When a job is submitted, the query is searched against this database with `nhmmer` (Wheeler and Eddy, 2013). Hits with *E ≤* 1 are listed, keeping the best hit per chain, and hits with *E >* 0.01 are shown faded to mark them as weaker matches. Each hit is linked to its RCSB entry and shown with the entry title. It is also aligned to the query in Rfam model coordinates, with the residues that differ from the query highlighted. Users can therefore see at once whether a solved structure exists for their RNA or a close homolog. Such a structure can be used to validate the consensus secondary structure inferred from covariation.

To inspect alignments directly in the browser, we developed a lightweight interactive alignment viewer in plain JavaScript, with no external dependencies. It displays the input, Rfam seed and final alignments. By default, residues are colored by conservation following the R2R convention (Weinberg and Breaker, 2011): red when *≥*97% of sequences share the base, black for *≥*90% and grey for *≥*75%. The alignment can also be colored by nucleotide or by percent identity to the column consensus, as in Jalview (Waterhouse et al., 2009). Above the alignment, the viewer shows a ruler, the consensus sequence, a percent-identity profile and an R2R-style row of circles that marks how often each column is occupied rather than gapped. Each sequence is annotated with its percent identity to the consensus, and sequences can be sorted by this value. Below the alignment, the consensus secondary structure predicted by CaCoFold (Rivas, 2020) is shown, and the base pairs with significant covariation according to R-scape (Rivas et al., 2017) are highlighted in green across the whole alignment. Hovering over a column shows its position, consensus nucleotide, conservation and occupancy.

Consensus secondary structures can be explored interactively with RFview, the RNA Framework structure viewer (Incarnato et al., 2018). RFview is shown alongside the static R2R drawings (Weinberg and Breaker, 2011) both displaying the R-scape/CaCoFold results (Rivas et al., 2017). It supports radiate and NAView layouts, zooming and coloring by alignment features. Finally, an interactive alignment viewer displays the input, Rfam seed and final alignments. Columns are colored by conservation following the R2R convention, and the base pairs that R-scape finds significantly covarying are highlighted. The viewer is accompanied by alignment summary statistics: number of sequences, effective number of sequences and percentage of canonical base pairs. Together, these tools let users judge the quality of an alignment and the evidence for a conserved structure directly in the browser, without downloading files or using external software.

### Addition of specialized databases

In addition to the main taxonomic genome databases, RNAhub and RNAhub2 now offer specialized genome collections for organelles, under-represented eukaryotic lineages and viruses. For eukaryotes and organelles, users can search all 327 Euglenozoa genomes available at NCBI and 15,233 NCBI RefSeq chloroplast and other plastid genomes (O’Leary et al., 2016). The Genome Taxonomy Database (GTDB) provides 901,341 bacterial and archaeal genomes from both isolates and metagenomes (Parks et al., 2022), extending well beyond the genomes of cultivated species. Viruses are covered by several complementary collections. These are 130,269 NCBI RNA virus and retrovirus genomes (Riboviria), and about 378,000 RNA virus sequences assembled from environmental metatranscriptomes (RVMT, RiboV1.4) (Neri et al., 2022). They also include the IMG/VR database of cultivated and uncultivated viral sequences from metagenomes (Camargo et al., 2023), and INPHARED2, a curated collection of 41,611 complete bacteriophage genomes from the Millard Lab (Cook et al., 2021). These collections let users search for structured RNAs in organelles, in lineages that are poorly represented in the main databases, and across RNA and DNA viruses and the uncultivated microbial diversity found in metagenomes.

## METHODS

### Query-adapted genome database construction

In the RNAhub2 workflow, each query is first used to perform a single homology broad search against the selected genome database. Candidate regions recovered during this prefiltering step are then extracted and concatenated to generate a reduced, query-specific search database, which is used as input for the standard RNAhub iterative alignment-building procedure with nhmmer (Wheeler and Eddy, 2013) and structural evaluation with R-scape (Rivas et al., 2017).

For a given input sequence, RNAhub2 performs independent homology searches against each NAF-compressed whole genome using nhmmer (Wheeler and Eddy, 2013) with a lenient E-value cutoff of 10.0, via the command `unnaf --fasta <genome.naf> | nhmmer --cpu 1 --dna --incE 10 <input sequence> -`. The nhmmer output alignments of homologs from all genomes are then unaligned and combined to form the query-dependent database, which is used for all subsequent homology searches.

Information about each sequence homolog in the query-adapted candidate database is summarized in an output table, where candidate sequence names, genomic coordinates, strand information, scores, and E-values are reported. This table provides a transparent record of the query-adapted database construction step and can also be used to apply additional E-value-based filters to the candidate set without rerunning the full genome-wide prefiltering search.

RNAhub2, like the original RNAhub, also supports searches in “flank mode”, in which the input sequence is extended with adjacent, potentially more conserved regions (for instance, an intron flanked by exon sequence). A preliminary alignment is first constructed using homologs of the extended (flanked) sequence and is then trimmed to represent only the query region. For RNAhub2 searches performed in flank mode (Gao et al., 2021), the query-adapted database is constructed using the entire query sequence, including flanks. The resulting candidate database is then passed to the standard RNAhub flank-mode workflow, in which the query-containing region is eventually extracted and processed as in the original RNAhub pipeline.

### Taxonomic information for genomes included in the RNAhub2 databases

The RNAhub genome databases were initially annotated only with NCBI GenBank assembly accessions (`GCA` identifiers). To assign taxonomic context to each genome, assembly accessions from the RNAhub2 collection were queried using the NCBI Datasets command-line tool (O’Leary et al., 2024), as follows. First, `datasets summary genome accession` was used to retrieve the corresponding NCBI Taxonomy ID and organism name for each assembly. The resulting TaxID was then used as input to `datasets summary taxonomy taxon`, which provides the full taxonomic classification. Kingdom, phylum, class, order, and family-level information was extracted using `jq` and merged into a single taxonomy table for each RNAhub2 database.

This procedure was applied independently to the Vertebrates, Invertebrates, Fungi, and Plants genome databases, producing a taxonomy table for each. These tables provided the taxonomic context required to summarize database composition and coverage. In addition, genomes were organized into nested, taxonomy-based directory structures, with genome files linked into folders corresponding to successive taxonomic levels. This organization enables database inspection and downstream analyses at different levels of taxonomic resolution.

The taxonomy-annotation pipeline is available as a GitHub repository https://github.com/ amkilar/fetch_taxonomy.

### Selection of extended RNAhub2 genome databases

Taxonomic representation of selected RNAhub genome databases was evaluated by combining genome selection lists, taxonomy tables, and genome-size estimates.

For each database, selected genomes were determined from RNase MRP-based subclustering [method developed in (Magnus et al., 2025)], using one representative per cluster at specific thresholds (0.60 for plants, 0.80 for vertebrates, 0.60 for invertebrates, 0.70 for fungi, and 0.825 for mammals). Genome accessions from these sets were matched to taxonomy and genome-size tables, with accessions serving as identifiers. For each taxonomic group, we calculated the total number of available genomes, the number of genomes included in the RNAhub database, and the percentage of available genomes represented in RNAhub. To estimate how many additional genomes could be included in the updated RNAhub databases after genome compression, we used precomputed clustering outputs, retaining one representative genome accession per cluster at each increasing thresholds. These accession lists were matched to genome metadata tables containing genome accession identifiers, compressed genome sizes, and taxonomic annotations. For each clustering threshold, the total number of retained genomes and their cumulative compressed size were calculated. These values were compared to the storage size of the original RNAhub database and to the size of the genome set selected with the previously used clustering threshold. This allowed us to identify new, higher clustering thresholds that increased the number of represented genomes, while keeping the total compressed database size within the storage quota originally allocated for each taxonomic category database.

### Benchmark of RNAhub and RNAhub2 on eight *C. elegans* non-coding RNAs

Eight annotated non-coding RNA genes from *Caenorhabditis elegans* were extracted as a FASTA sequence from Rfam: tRNA (BX284606.5–3994608-3994666), 5S rRNA (BX284605.5–17118958-17118837), U4 (BX284605.5–11391214-11391076), U1 (BX284602.5–12944967-12944804), a C/D-box snoRNA (BX284603.4–502803-502889), U2 (BX284602.5–6716354-6716168), RNase P RNA (BX284602.5– 7200233-7200415), and SSU rRNA (BX284601.5–15062084-15063836).

Two invertebrate genome databases were used, referred to by the RNase MRP identity threshold at which genomes were subclustered: Invertebrates at 60% clustering threshold (invertebrate-60) (165 genomes, 14 of them Nematoda) and Invertebrates at 80% clustering threshold (invertebrate-80) (1,895 genomes, 104 Nematoda). Each query was run through the original RNAhub pipeline (Magnus et al., 2025) against invertebrate-60 and through RNAhub2 against invertebrate-60, so that both pipelines were run against the same database, and then through RNAhub2 against invertebrate-80 (Table 2).

For each run we recorded the total wall-clock runtime, from query submission to completion of the R-scape/CaCoFold evaluation. From the `v last stats.txt` output file we parsed the number of aligned sequences, the average pairwise percent identity, and the average aligned sequence length; from the `cacofold.power` output file, the number of base pairs observed to covary at the R-scape significance threshold.

There were two runs that had to be handled individually. For the tRNA query under the original RNAhub, the first-iteration E-value threshold was relaxed from 1 *×* 10*^−^*^10^ to 1 *×* 10*^−^*^5^, matching the value used in the remaining iterations; RNAhub2 was run on this query with the standard workflow. The SSU query did not complete against invertebrate-80, where prefiltering recovered 165,428 candidate sequences and the intermediate databases reached 11–12 GB; This is expected given the high number of copies of SSU in eukaryotic genome, including pseudogenes. SSU is reported for invertebrate-60 only.

### Comparison of RNAhub and RNAhub2 using selected CASP RNA targets

We compared RNAhub and RNAhub2 using 14 selected RNA targets. These comprised 11 CASP16 targets representing established structural RNA families (Table 3) and three eukaryotic targets: the CASP16 group I intron R1260, the CASP15 CPEB3 ribozyme R1107, and the CASP17 U1 snRNA R2415s0 (Fig. 5). The comparisons assessed homolog recruitment and recovery of significant covariation.

The 11 CASP16 targets in Table 3 were analyzed in unflanked mode using the database combinations specified in the table. Searches used the Viral or Bacteria databases, or combinations including GTDB and IMG/VR. The bacterial and viral genome collections were unchanged between RNAhub and RNAhub2.

For the three eukaryotic targets, RNAhub searches used the original genome databases and RNAhub2 searches used the corresponding expanded collections. R1260 was searched against Invertebrates; R1107 against the combined Mammals and Vertebrates databases; and R2415s0 against the combined Mammals, Vertebrates, Invertebrates, Fungi, and Plants databases.

For Table 3, we recorded the number of aligned sequences, mean pairwise sequence identity, and number of significantly covarying base pairs for the final nhmmer alignment (`v3 rm`) and, where available, the Infernal realignment. Covariation statistics were obtained from the R-scape `cacofold.power` output, using a significance threshold of *E <* 0.05. Pairs classified by R-scape as possibly arising from protein-coding constraints were reported separately in parentheses. For Figure 5, we additionally recorded alignment length, mean sequence length, the number of base pairs in the CaCoFold model, and the expected number of covarying base pairs. Figure statistics correspond to the Infernal realignment where available. Neither R1107 analysis detected significant covariation, and Infernal realignment was not performed. The original RNAhub search for R1260 terminated after the first iteration without recruiting homologs.

### Standalone workflow for RNAhub2 and RNAhub

In addition to the web server, RNAhub and RNAhub2 can be run from the command line using the user’s own computational resources. Standalone execution requires users to supply their own genome databases. The curated genome collections maintained by RNAhub are not distributed with the software. Users wishing to search these curated collections should use the web server at https://rnahub.org.

Both workflows are accessible through `rnahub.py`, with the query supplied through `--input` and the search database through `--db`. By default, the program runs the original RNAhub workflow, performing iterative nhmmer searches against user-supplied FASTA databases, either uncompressed or gzip-compressed. Adding `--minidb` enables the RNAhub2 workflow. In this mode, `--db` specifies directories containing individual genomes in NAF-compressed or uncompressed FASTA format (`.fa`, `.fasta`, or `.fna`). The program first searches these genomes to construct a query-adapted candidate database, which is used for subsequent iterative searches. NAF input is supported only with `--minidb`. Genomes in this format must otherwise be decompressed before analysis.

Unflanked analysis is the default in both workflows and requires no additional flag. Flanked analysis is enabled by supplying an extended query sequence through `--flanked`, alongside the query region supplied through `--input`. Alternatively, `--flanks-in-header` identifies the query region within an extended input sequence using coordinates encoded in its FASTA header. In flanked mode, the extended sequence guides homolog recruitment before the alignment is trimmed to the query region. Both workflows provide downstream structural assessment with R-scape/CaCoFold and optional Infernal realignment. Source code and installation instructions are available at https://github.com/mmagnus/rnahub.

## DISCUSSION

RNAhub2 addresses the two main limitations of the original RNAhub webserver. The query-adapted prefiltering restricts genome-wide homology searches to a reduced, query-specific database before the iterative alignment building step, substantially reducing runtime for searches against large eukaryotic databases. Independently, NAF compression reduces genome storage by approximately 4.2-fold, which we used to expand the RNAhub2 databases by nearly 5,000 additional eukaryotic species and to improve taxonomic balance within existing databases, all within the original storage quota. Genome compression and query-adapted prefiltering therefore make complementary contributions: compression allows more genomes to be stored, while prefiltering reduces the time required to search them iteratively. Together, these improvements make the additional genome diversity more accessible for comparative structural analysis. Expanding taxonomic sampling can substantially strengthen covariation signal, as illustrated by the human telomerase RNA, where the expanded vertebrate database increased the final alignment from 56 to 290 sequences and provided structural support for four stems, while RNAhub provided support for only one stem. However, this benefit is not automatic: it depends on whether the added sequences carry informative substitutions, rather than being redundant or overly divergent homologs (Rivas et al., 2020). Our benchmark on *C. elegans* ncRNAs demonstrates the overall benefit of expanding the invertebrate database. For RNase P and a C/D-box snoRNA, the larger database increased alignment depth and generally improved covariation power, despite a decrease in pairwise identity. The number of significantly covarying base pairs for 5S rRNA decreased slightly, but this represents an exception rather than a general trend; such small differences can arise because individual base pairs may lie close to the R-scape significance threshold.

The eukaryotic CASP targets further illustrate that the effect of database expansion depends on the evolutionary information contributed by the additional sequences. For R1260, homologs available in the expanded invertebrates database provided the initial sequences needed to construct an informative search profile and recover covariation-supported structural features by RNAhub2, which was not the case for RNAhub. However, while RNAhub2 produced substantially deeper alignments for R1107 and R2415s0, this did not increase the number of significantly covarying pairs. The extended RNAhub2 genome database therefore increases the opportunity to recruit informative homologs, but greater alignment depth alone does not guarantee stronger covariation support. The outcome also depends on the evolutionary variation, homology, and alignment quality of the recruited sequences.

The bacterial and viral CASP16 comparisons highlight a separate trade-off inherent in restricting iterative searches to a query-adapted database. Although this approach reduces computational cost, it makes subsequent homolog recruitment dependent on the initial query: sequences excluded during prefiltering remain unavailable even as the search profile improves through iteration. This provides a possible explanation for target-dependent losses of structural signal, although its contribution remains to be established by tracing recruitment through the workflow. An important distinction is therefore between insufficient comparative evidence produced by a particular search and evidence against a conserved structure. Searches yielding little or no covariation should be interpreted in the context of alignment quality and statistical power.

The current implementation also has practical limits. Although prefiltering accelerates the iterative searches, the initial search against the selected genome collection can still account for most of the total runtime. A separate limitation arises for queries with many genomic matches, such as the SSU rRNA benchmarked here. These queries can generate candidate databases so large that the subsequent iterative searches exceed the available memory, indicating that further optimization may be needed for highly conserved RNAs present in many genomic copies. In addition, although compression substantially increased the number of genomes that fit within our storage quota, the eukaryotic databases remain a subsample of the full diversity of genome assemblies now publicly available; further gains in storage or compression efficiency would allow additional taxonomic expansion.

Overall, combining query-adapted prefiltering with compressed, taxonomically balanced genome databases makes RNAhub2 substantially faster than RNAhub while preserving, and in many cases improving, the sensitivity of covariation-based structure discovery, allowing eukaryotic comparative RNA searches to draw on substantially larger and more taxonomically diverse genome collections than were previously practical within the same runtime and storage quota. The improvements included in RNAhub2 will therefore facilitate discovery of structural ncRNAs across large eukaryotic genome.

## AUTHOR CONTRIBUTIONS

E.R. and Q.V. conceived the general idea; A.M.K. and W.G. developed the RNAhub2 workflow. M.M. and A.M.K. updated the RNAhub webserver. A.M.K., W.G., N.D., and M.M. performed experiments. A.M.K. and E.R. wrote the manuscript. All authors edited the manuscript. All authors have read and agreed to the published version of the manuscript.

## DATA AVAILABILITY

The RNAhub2 workflow is available at https://rnahub.org. This website is free and open to all users without a login requirement. The source code and installation instructions are available at https://github.com/mmagnus/rnahub.

## FUNDING

This work was supported by the US National Institutes of Health (NIH) grant 1R21GM148902-01 to E.R. and Q.V. M.M. was supported by the Polish National Science Centre (NCN SONATA BIS, grant 2025/58/E/NZ1/00373 to M.M.). For the purpose of open access, the authors have applied a CC BY public copyright license to any Author Accepted Manuscript version arising from this submission.

## CONFLICTS OF INTEREST

The authors declare no conflict of interest.

---

**Supplementary Table S1.** Alignment statistics for selected eukaryotic RNA sequences analyzed in unflanked mode. Statistics for the final nhmmer alignment (v3 rm) are shown in black (top), and those for the Infernal realignment are shown in blue (bottom). Numbers in parentheses indicate significantly covarying pairs classified by R-scape as possibly arising from protein-coding constraints.

| Sequences<br>Rfam ID | Database | Pairwise ID |  | #sequences |  | #covarying pairs |  |
|---|---|---|---|---|---|---|---|
|  |  | rnahub | rnahub2 | rnahub | rnahub2 | rnahub | rnahub2 |
| Fungal U3 | Fungi | 61.94%<br>61.95% | 70.69%<br>70.48% | 1953<br>1953 | 2623<br>2623 | 71(32c)<br>73(26c) | 20(5c)<br>31(5c) |
| Fungal SRP | Fungi | 59.54%<br>58.69% | 63.81%<br>60.58% | 155<br>155 | 116<br>116 | 24(8c)<br>35(2c) | 8<br>28 |
| Plant U3 | Plants | 69.48%<br>68.59% | 72.68%<br>72.57% | 74<br>74 | 1045<br>1045 | 4<br>8 | 7<br>10 |
| Plant SRP | Plants | 63.82%<br>63.62% | 66.15%<br>65.96% | 73<br>73 | 1044<br>1044 | 1<br>12 | 15<br>43 |
| Invertebrate U3 | Invertebrate | 94.842%<br>NA | 63.86%<br>63.26% | 3<br>NA | 1199<br>1199 | 0<br>NA | 3(1c)<br>4(1c) |
| Invertebrate SRP | Invertebrate | 65.73%<br>65.59% | 69.29%<br>69.46% | 163<br>163 | 1876<br>1876 | 35<br>62 | 46<br>73 |

## SUPPLEMENTARY MATERIAL

### NAF compression substantially reduces genome storage without increasing search time

One of the main practical challenges in extending RNAhub to larger taxonomic databases is the storage requirement associated with whole-genome sequence collections. This limitation is particularly important for plant genomes, where both the number of available assemblies and the broad variation in genome size make it difficult to include all candidate genomes within the storage limits of the web server. To evaluate whether compressed genome representations could reduce storage requirements while preserving acceptable search performance, we benchmarked several compression formats on a representative set of plant genomes.

This benchmark was designed to approximate the RNAhub2 use case. In this setting, genomes are stored in compressed form to reduce disk usage and decompressed only at query time. The comparison therefore evaluates the combined cost of decompression and nhmmer search, rather than decompression time alone. This distinction is important, as compression is performed only once during database preparation, whereas decompression and search are carreid out for each user query.

The genome compression benchmark shows that compressed storage can substantially reduce the disk space required for large genome databases (Fig. S1, A). This is particularly relevant for plant databases, where inclusion of all available representative genomes is limited by the storage quota of the web server. Moreover, the benchmark included genomes ranging from approximately 1.3 Gb to more than 23 Gb, thereby testing compression behavior across both moderate and very large plant assemblies.

Among the tested compression formats, NAF provides the most favorable balance between storage reduction and search performance (Fig. S1, B). NAF-compressed genomes are approximately 4.5-fold smaller than the corresponding uncompressed FASTA files, while the total runtime of decompression followed by nhmmer search remains comparable to the uncompressed FASTA control. In contrast, some general-purpose compression methods show less favorable trade-offs. For example, lz4 provides only modest compression and increased relative computation time, whereas xz compression achieves strong compression but shows more variable behavior across genome classes. These results indicate that compression ratio alone is not sufficient for selecting a storage format for RNAhub2 databases.

The advantage of NAF compression is consistent across all tested genome-size classes. Small genomes, repeat-rich grass genomes, medium-sized genomes, and very large genomes all show substantial compression relative to FASTA, without a corresponding increase in nhmmer search time. This suggests that NAF-compressed genome storage is an effective way to increase database coverage while preserving the interactive search performance required in a web-server environment. These results support the use of NAF as the preferred storage format for the expanded RNAhub2 genome databases.

**Supplementary Figure S1.**
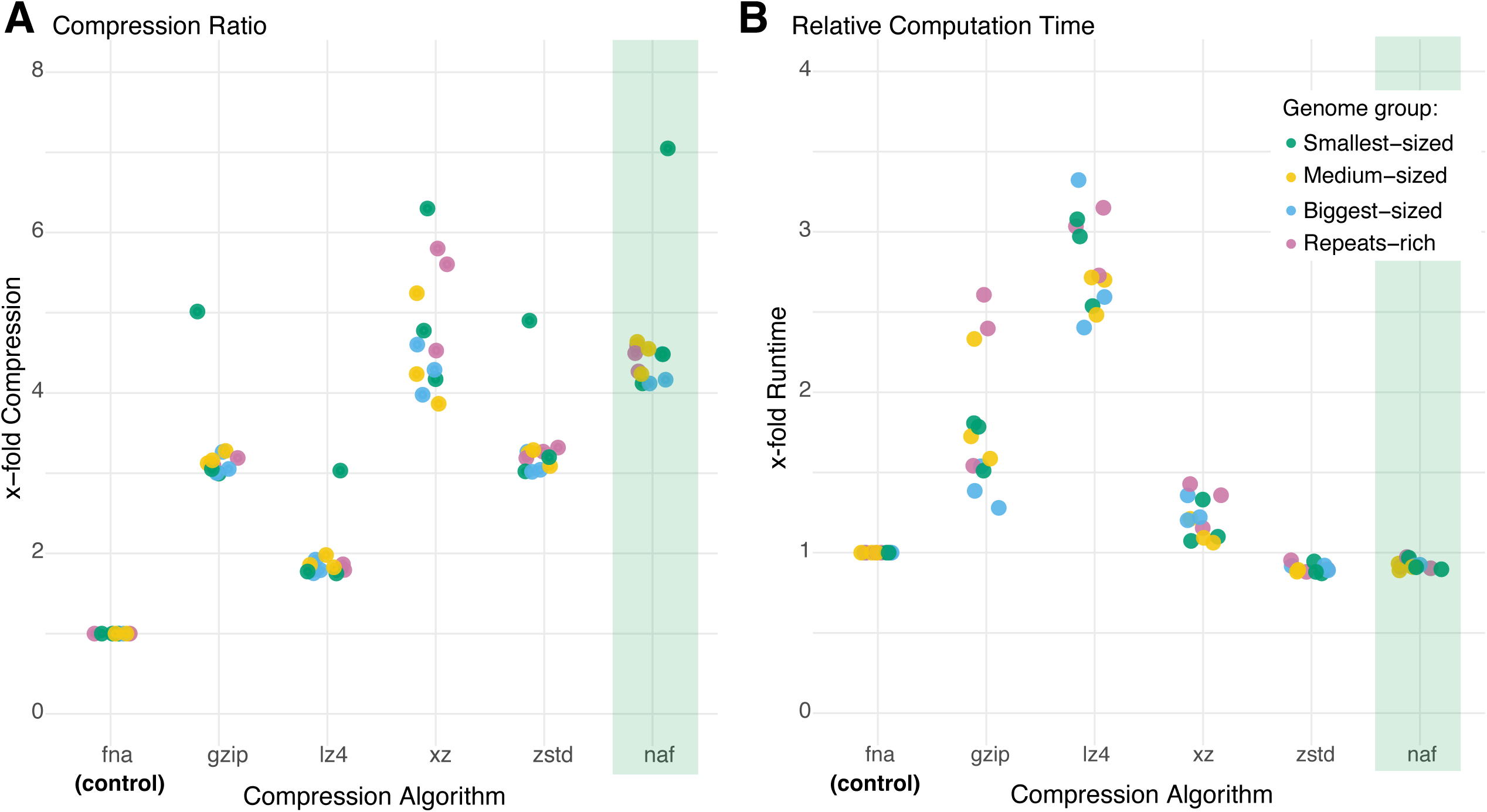
Effect of genome compression format on storage reduction and search runtime. Comparison of six genome storage formats for a representative plant genome set: uncompressed FASTA (fna), gzip, lz4, xz, zstd, and naf. (A) Compression ratio relative to uncompressed FASTA, with higher values indicating stronger storage reduction. (B) Relative computation time compared to uncompressed FASTA, measured as the combined decompression and nhmmer search runtime, with FASTA normalized to 1. Each point represents one genome, colored by genome group: smallest-sized, medium-sized, biggest-sized, or repeat-rich. The green bar indicates the compression algorithm selected for RNAhub database storage.

### Compression benchmarking using plant genomes

To evaluate genome compression effect on storage and runtime, we selected a set of plant genomes with a broad range of genome sizes, from approximately 1.3 Gb to more than 23 Gb. We also included genomes of similar size but with different repeat composition, including both moderate-sized angiosperm genomes and very large gymnosperm and fern genomes.

The small-genome set includes *Erythroxylum citrifolium* (`GCA 026284465.1`; 1303 Mb), *Warburgia ugandensis* (`GCA 035236585.1`; 1305 Mb), *Phytolacca americana* (`GCA 040581565.1`; 1307 Mb), *Fagopyrum esculentum* (`GCA 033239045.1`; 1472 Mb), *Poa infirma* (`GCA 040144145.1`; 1339 Mb), and *Bothriochloa decipiens* (`GCA 023333625.1`; 1412 Mb). The grass genomes are considered separately because their repeat-rich composition could influence compression efficiency and search performance.

The medium-to-large genome set include *Dryopteris marginalis* (`GCA 027576515.1`; 1413 Mb), *Rumohra adiantiformis* (`GCA 033807815.1`; 3229 Mb), and *Cibotium barometz* (`GCA 041138505.1`; 4054 Mb). The very large genome set include *Pseudotsuga menziesii* (`GCA 001517045.1`; 17188 Mb), *Torreya grandis* (`GCA 037957365.1`; 22130 Mb), and *Picea mariana* (`GCA 032191745.1`; 23098 Mb).

For each genome, the uncompressed FASTA file is used as the reference input. The FASTA files are compressed using `gzip` (`gzip -5 -c`), `lz4` (`lz4 -c`), `xz` (`xz -c`), `zstd` (`zstd`), and NAF (`ennaf`). Compression ratios are calculated as the uncompressed FASTA file size divided by the corresponding compressed file size, so that higher values indicate stronger storage reduction.

To assess whether compressed genomes could be used directly in the RNAhub2 search workflow, each compressed file was decompressed on-the-fly and streamed to nhmmer through a UNIX pipe, without creating intermediate FASTA files. Searches against uncompressed FASTA files serve as runtime reference. The benchmark measures the combined cost of decompression and nhmmer search, reflecting the expected web-server use case in which genomes are stored in compressed form and decompressed only during query execution.

We used *Arabidopsis thaliana* RNase MRP sequence (`URS00006B077B`) as query for all benchmarking searches. Runtime ratios were calculated as the runtime for each compressed-input search divided by the runtime obtained using the corresponding uncompressed FASTA file. The results of the benchmark are reported in Supplemental Fig. S1.

## Notes

https://rnahub.org/

### Competing Interest Statement

The authors have declared no competing interest.

### Summary of Updates

The pdf uploaded before did not display the references correctly. This version includes the references correctly

